# Heterogeneous folding landscapes and predetermined breaking points within a protein family

**DOI:** 10.1101/2024.04.22.590563

**Authors:** Sebastian Pechmann

## Abstract

The accurate prediction of protein structures with artificial intelligence has been a spectacular success. Yet, how proteins fold into their native structures inside the cell remains incompletely understood. Of particular interest is to rationalize how proteins interact with the protein homeostasis network, an organism specific set of protein folding and quality control enzymes. Failure of protein homeostasis leads to widespread misfolding and aggregation, and thus neurodegeneration. Here, I present a comparative analysis of the folding across a protein family from a single organism, the *Saccharomyces cerevisiae* small GTPases. Using computational modelling to directly probe folding dynamics, this work shows how near identical structures from the same folding environment exhibit heterogeneous folding landscapes. Remarkably, yeast small GTPases are found to unfold along different pathways either via the N- or C-terminus initiated by structure-encoded predetermined breaking points. Moreover, degrons as recognition signals for ubiquitin-dependent degradation were systematically depleted from the initial unfolding sites, as if to protect from too rapid degradation upon spontaneous unfolding. In turn, Hsp70 Ssb chaperone binding correlated only with N-terminal hydrophobicity. The presented results highlight a direct coordination of folding pathway and protein homeostasis interaction signals across a protein family. A deeper understanding of the interdependence of proteins with their folding environment will help to rationalize and combat disease linked to protein misfolding and dysregulation. More generally, this work underlines the importance of understanding protein folding in the cellular context, and highlights valuable constraints towards a systems-level understanding of protein homeostasis.

## Introduction

The highly accurate prediction of protein structures with algorithms based on artificial intelligence is revolutionizing protein science [1]. Readily available structural models for entire proteomes are impacting essentially all aspects of biological research, with some of the most impressive successes so far thus achieved in the engineering of proteins with new or improved properties [2, 3]. However, proteins are synthesized by ribosomes as linear chains of amino acids that generally have to first fold, inside the crowded cell, into their characteristic three-dimensional structures before they become active. The question of how proteins fold remains an incompletely understood problem [4] that is not solved by current abilities to predict protein structures [5, 6].

Importantly, proteins are not generally optimized for structural stability alone [7], but comprise complex biomolecules under myriad different and partly opposing constraints [8]. Foremost, proteins do not exist in static shapes but rather as dynamic ensembles of structures [9]. Often, the allosteric motion between such different conformations is prerequisite for a protein’s function [10, 11]. Consequently, protein folding landscapes involve inherent trade-offs between protein stability and function [12, 13], and thus between function and folding [14] of proteins that are generally only meta-stable [15] and marginally soluble [16]. Small proteins can nonetheless fold within milliseconds [17]. Longer proteins however bear an increased likelihood of forming non-native contacts [18] as well as often depend on multiple independent pathways [19, 20] and multiple intermediate states [21] for successful folding. In the extreme, different folding pathways can even lead to alternate final structures [22].

Furthermore, proteins fold inside the cell in tight interaction with and dependence on their environment. An elaborate regulatory network composed predominantly of protein folding [23] and quality control [24] enzymes, the protein homeostasis network [25], monitors and assists the whole protein lifecycle from synthesis and folding to translocation and degradation. In eukaryotes, the vast majority of proteins interacts with molecular chaperones [26], which can fundamentally alter folding landscapes. However, central details regarding chaperone interactions including their selectivity remain unclear. Moreover, unlike the protein amino acid sequences that compare across taxa, chaperone networks and thus the cellular folding milieus can vary dramatically across species [27]. To this end, understanding how proteins fold inside the cell in interaction with the protein homeostasis network is not only of paramount importance for understanding protein folding itself, but also forms the basis for the regulation of most aspects of cell (dys)function [28], such as signalling [29], synaptic plasticity in neurons [30], or development of cancer [31].

One elegant and controlled approach to inform on the interplay of protein folding and homeostasis builds upon comparative analyses of structural homologues within an organism. For instance, the comparison of a set of *S.cerevisiae* proteins that fold into near identical structures but some with and some without dependence on the cotranslationally acting Hsp70 chaperones Ssb1/2 highlighted surprising insights into the determinants of these chaperone interactions [32]. Specifically, unlike long multidomain proteins with complex topologies that interact with Ssb to avoid misfolding and aggregation during cotranslational folding [33], the set of small single-domain proteins showed almost no correlation between chaperone dependence and folding challenges [32]. Instead, the presence of short recognition sequences was by far the strongest determinant of chaperone interaction [32], as if these were programmed into the sequences for other reasons such as regulatory fidelity. Taken together, many open questions regarding protein folding dynamics cannot be systematically answered from the analysis of static protein sequences and structures alone.

Here, bioinformatic analyses are combined with molecular modelling to explicitly probe protein equilibrium and (un)folding dynamics across a protein family. The comparison of 16 *S.cerevisiae* small GTPases from the G-Protein Family provides an ideal test case for uncovering general and differentiating characteristics of their folding landscapes. At the same time, the analysed small GTPases comprise a class of very important enzymes involved in protein sorting and trafficking [34] that are highly conserved [35], potential drug targets [36], and thus of high interest in their own right. Strikingly, this work shows how yeast GTPases differentially unfold from predetermined breaking points that are encoded in the native protein structures and that control the exposure of degrons as recognition sites of ubiquitin-dependent degradation.

## Results

Structurally homologous single-domain proteins from the same folding environment provide a uniquely controlled test case for studying similarities and constraints in protein folding landscapes. This rang particularly true for the 16 *S.cerevisiae* proteins from the family of small GTPases that aligned with exceptionally low backbone root mean squared deviation (Figure 1A) [32]. The characteristic GTPase fold under investigation consisted of six *β*-strands of which b2 is antiparallel to all others, and five flanking *α*-helices (Figure 1A). Even in absence of any conservation of site-specific contacts between the individual protein structures [32], these proteins comprised a well-defined core together with strong interconnectivity between the conserved and aligned secondary structure elements (Figure 1B).

**Fig. 1.**
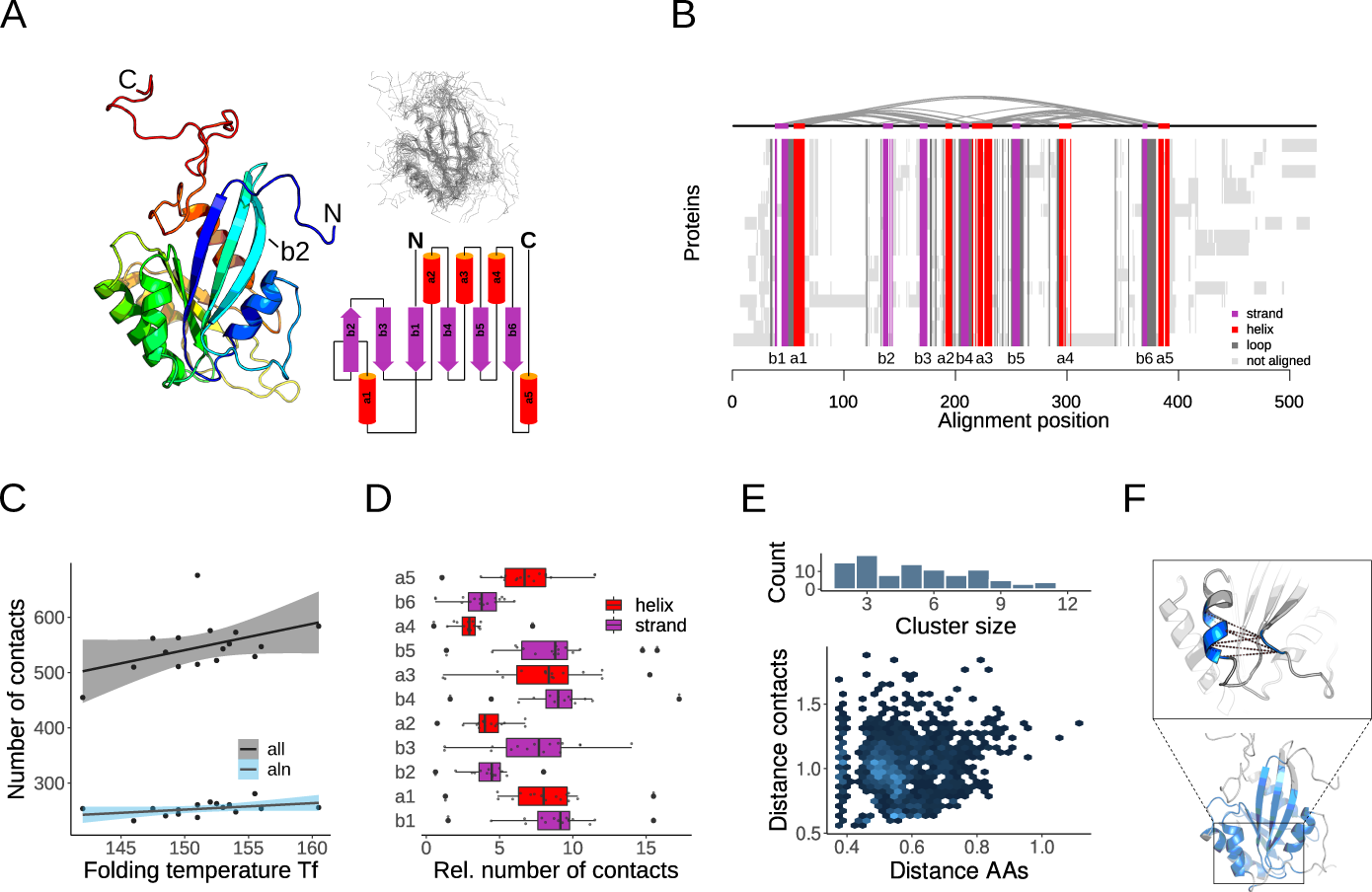
Native contacts and contact clusters in the *S.cerevisiae* protein family of small GTPases. **A** Secondary structure representation of the exemplary GTPase YPT53, shown together with the backbone alignment of all 16 homologous structures as well as a secondary structure diagram of their GTPase fold. **B** The structure-based multiple sequence alignment highlights tightly conserved secondary structure elements that are connected by a network of native contacts. **C** The number of native contacts in the different protein structures is shown as function of their corresponding folding temperatures *T_f_*. Also shown is the correlation between *T_f_* and the numbers of native contacts limited to only aligned positions. **D** Distributions of the per protein numbers of contacts in the different secondary structure elements. **E** Contact clusters provide a robust means to compare intramolecular distances even in absence of conserved individual contacts. Shown are the distributions of the numbers of contacts per contact clusters as well as the tight ranges of distances underlying their definition. Herein, the contact distance describes the contact length in nm between two residues, and the amino acid (AA) distance the proximity in nm between two contacts along the protein backbone. **F** Contact clusters cover 125 out of 128 aligned positions, thus are highly representative of the protein cores as shown by the coloured exemplary secondary structure representation where positions included in the contact clusters are shown in blue. The inlet highlights an exemplary contact cluster of 6 contacts.

### Contact clusters robustly compare distances across a protein family

I first determined the folding temperatures *T_f_* of the proteins, i.e. the temperatures at which the folded and unfolded states are equally likely, thus affording accurate and efficient sampling of folding dynamics. The total numbers of native contacts in the GTPases correlated with *T_f_*as expected (Figure 1C). An even stronger correlation could be observed when only considering the contacts in aligned regions (Figure 1C), which is characteristic for small single-domain proteins with a well-defined structural core. Of note, a range of different of *T_f_*values also indicated differences in thermo-stability. Grouped by secondary structure, the numbers of native contacts showed considerable variation (Figure 1D). Notably, the terminal secondary structure elements b1 and a5 exhibited broader distributions, thus differences between the proteins, while a2 and b2 were characterized by fewer contacts in narrower distributions (Figure 1D).

Next, to devise a set of characteristic distances that allow the comparison of folding dynamics in an unbiased and robust manner even without conservation of individual contacts across different proteins, I defined a set of contact clusters. This aimed to alleviate the discrepancies in the intramolecular distances between any two residues that were in direct contact in some proteins but not in the others. Specifically, the encompassing set of native contacts from all structures was mapped onto the alignment and clustered by proximity (*see Methods*). At a low cutoff of 1*nm*, this yielded 99 contact clusters that combined on average 5 distinct residue-residue contacts (Figure 1E). Because contacts within a cluster were close neighbours along the protein backbone, their average contact lengths provided strongly representative distances (Figure 1E, F). Importantly, 98% of the aligned positions were thus covered by contact clusters (Figure 1F).

### Global and local differences in frustration and flexibility

Before directly probing protein (un)folding, I sought to gain an overview of the energetics and dynamics in this protein family. Frustration in proteins quantifies how energetically favourably an amino acids contributes to the structure compared to all other possible amino acids, wherein higher scores indicate less favourable energetics [37]. The average frustration profile along the alignment revealed that the largest variation in the aligned regions of the proteins was found in helices a2 and a3 (Figure 2A). Overall, the helices a2-a5 were systematically more frustrated than the rest (Figure 2B). Thus, patterns of frustration across secondary structure elements were conserved even in absence of the conservation of individual contacts, as observed before [38]. Furthermore, the average frustration in some secondary structures varied considerably between proteins (Figure 2B). Similarly, while the observation of higher frustration in non-aligned regions by itself is trivial (Figures 2A,C), there were also noticeable global differences in the average frustration of the aligned and non-aligned regions of the different proteins (Figure 2C).

**Fig. 2.**
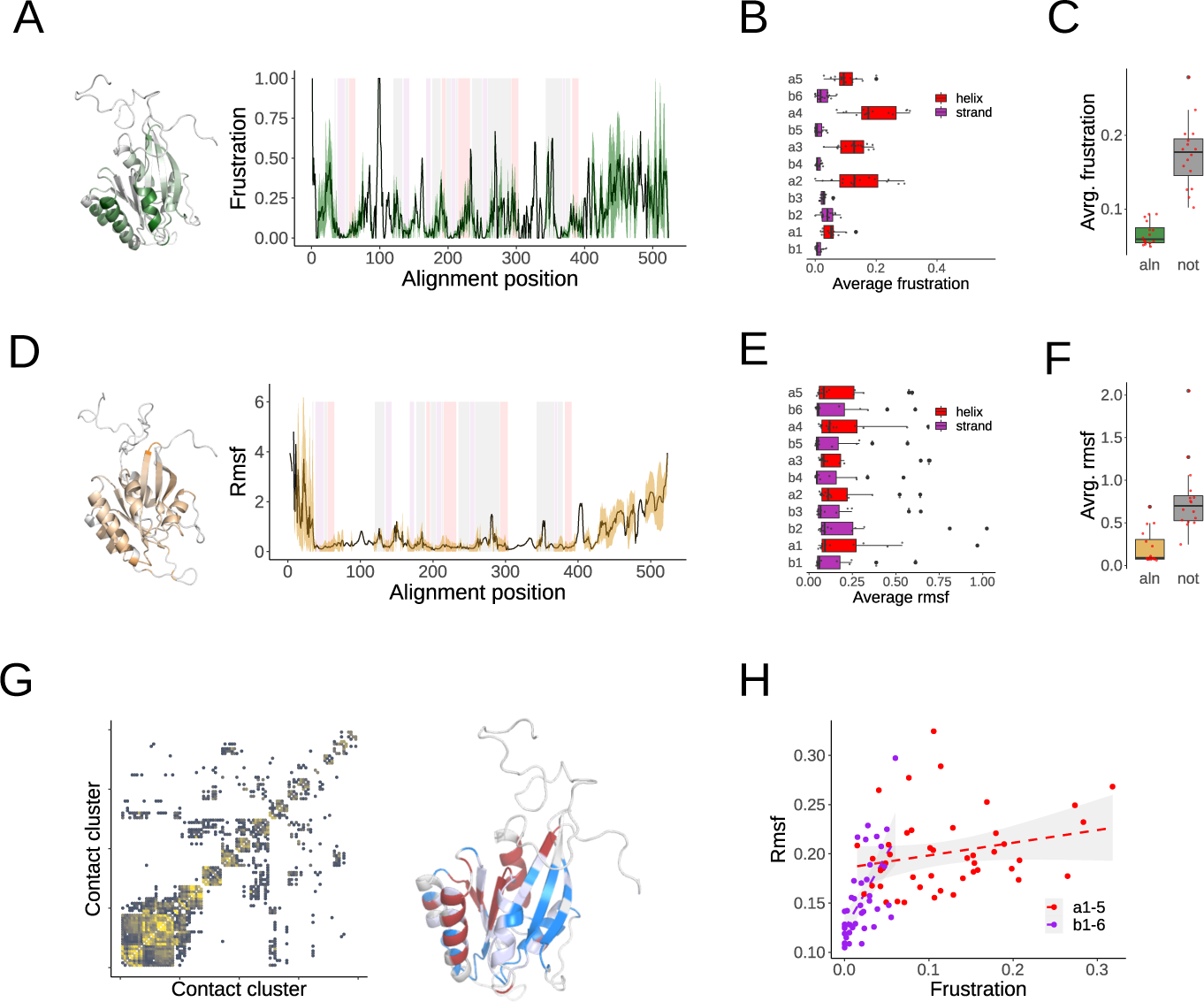
Frustration and flexibility in structural homologues. **A** The frustration profile represented by mean and standard-deviation across the alignment is shown together with the positions of the aligned secondary structures. The gradient-coloured exemplary structure highlights the positions with the most variation between the different proteins. **B** Distributions of average frustration scores for the individual secondary structures. **C** Distributions of average frustration scores for aligned and unaligned regions. **D** The profile of root mean squared fluctuations (rmsf) as in (A). The highest variable regions are highlighted on an exemplary structure. **E** Distributions of average rmsf values in secondary structures. **F** Distributions of average rmsf values for aligned and unaligned regions. **G** Map of correlated motions between contact clusters. Brighter color-coding indicates a higher number of individual proteins for which the distance profiles of two contact clusters is correlated. Consolidating the non-overlapping parts of the clusters of correlated motions divides the protein roughly into two halves of correlated motions, as shown on the structure in red and blue. **H** Correlation between frustration and protein flexibility (rmsf). Each data point represents the averages for one secondary structure element in one protein.

Local energetics influence protein stability and flexibility, which can similarly vary across a protein family [39]. Protein motions quantified through their root mean squared fluctuation (rmsf) in trajectories of molecular dynamics simulations (*see Methods*) were overall similar (Figures 2D,E). Noteworthy were consistently higher rmsf values in the helices, and in b2 compared to the other *β*-strands (Figure 2E). Moreover, some proteins exhibited overall higher rmsf values than others (Figure 2F). Next, I computed a map of correlated motions from the contact cluster distance profiles along the simulation trajectories (*see Methods*). Two regions of correlated motions that could systematically be detected across the protein family divided the protein roughly into an N-terminal half of *α*-helices a1/a2 and a *β*-sheet of b1-b3, as well as a C-terminal half of a3-a5 and b4-b6 (Figure 2G).

Importantly, for averages across secondary structure elements protein flexibilities and levels of frustration were directly correlated (Figure 2H). The lower frustration and motion, as well as higher sensitivity to changes in frustration in *β*-strands likely reflected their high propensity to form harmful protein aggregates [40]. In turn, the more dispersed correlation for *α*-helices pointed to regions of more pronounced differences between the different GTPases.

### Heterogeneous unfolding via predetermined breaking points

Protein topology strongly influences available folding pathways [41], but not sufficiently so to make them easily predictable. To directly probe protein folding dynamics, I thus next ran a large number of coarse-grained unfolding simulations (*see Methods*). These exploited the efficiency of structure-based or Gō models that only consider native contacts [42], and built upon the assumption, generally assumed valid for small single-domain proteins, that the unfolding pathway is the reverse of the folding pathway [43].

The time evolution of the intramolecular distances defined by the contact clusters allowed to robustly compare the unfolding of structural homologues (Figure 3A). Specifically, the systematic determination of the breakpoints of all contact cluster in each simulation trajectory provided a detailed description of the unfolding pathways (Figure 3B). Immediately noticeable was pervasive heterogeneity in how these proteins unfolded. To better understand these differences, I next examined the first breaking contact clusters in each simulation to map how the unfolding was initiated (*see Methods*). Strikingly, essentially all trajectories grouped into one of five classes of initial unfolding based on the location of these first breaking contact cluster (Figure 3C). The most common initiation of unfolding, observed in 43% of the simulations, was the breaking of a contact cluster between a2 and b2 (Figure 3C, D). The second most common mode of unfolding was via the loosening of the a5 *α*-helix near the C-terminus, followed unfolding via destabilization of contact clusters in a3 (Figure 3C, D). Two unfolding classes, “a1”, and “a2/b2” were on the N-terminal half of the protein while the unfolding classes “a3”, “a4/b5”, and “a5” involved the breakage of contact clusters in the C-terminal half.

**Fig. 3.**
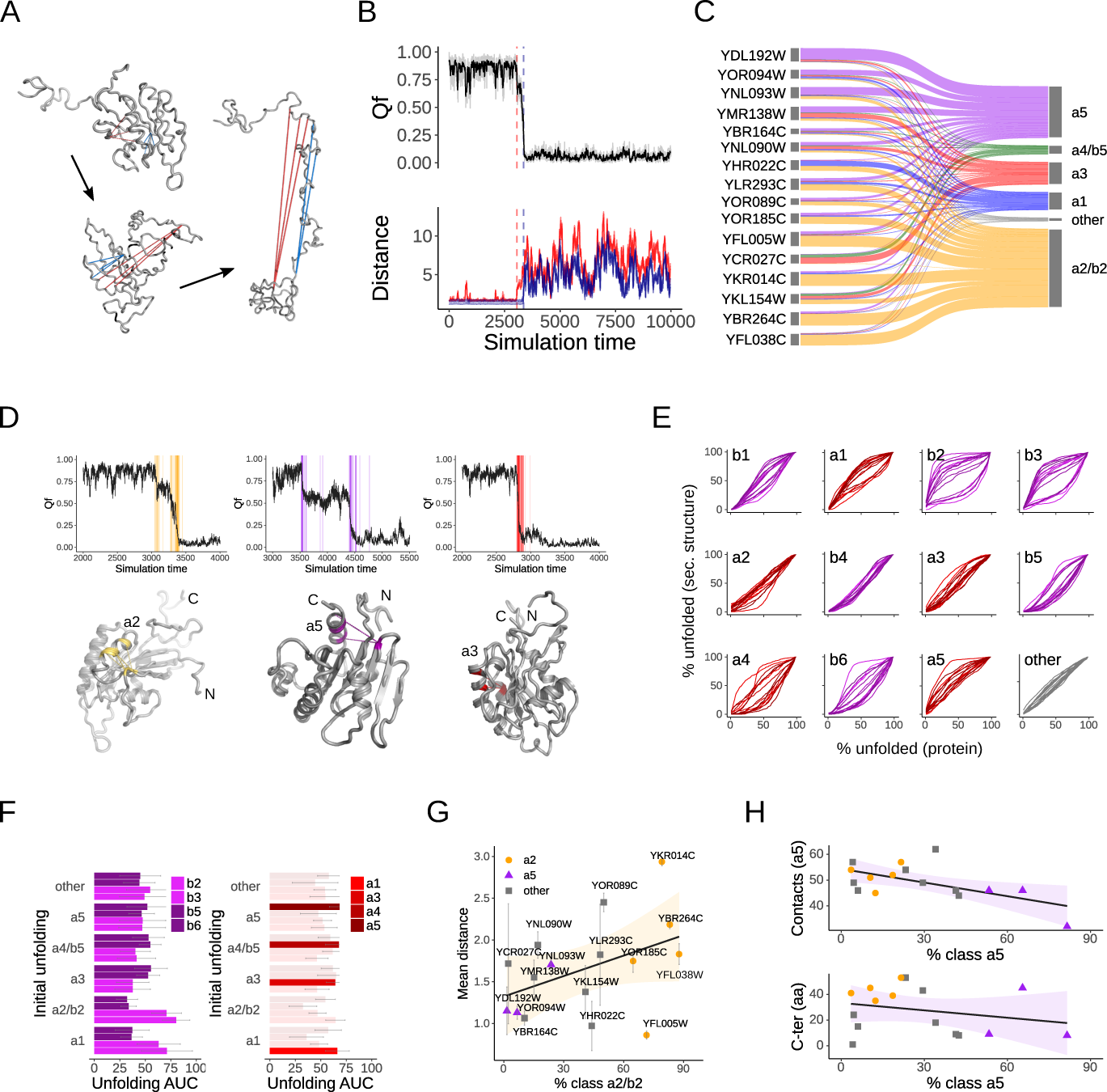
Unfolding via predetermined breaking points. **A** Exemplary illustration of three structural states along an unfolding trajectory. The contacts of two contact clusters are shown in red and blue respectively. **B** Exemplary profiles of an unfolding simulation. *Q_f_*quantifies the overall fraction of formed contacts and indicates the overall unfolding along the simulation trajectory. The distance profiles of all contacts from two exemplary contact cluster are shown in red and blue. Breakpoints within a contact cluster are highly synchronous, and shown with dashed lines. The determination of all contact cluster breakpoints during an unfolding simulation provides a detailed description of the unfolding pathway. **C** Classification of simulation trajectories into unfolding classes based on the location of first breaking contacts. The mapping illustrates the fraction of simulations for each protein that unfold via unfolding classes “a1”, “a2/b2”, “a3”, “a4/b5”, “a5” and “other”. **D** Exemplary unfolding trajectories and locations of the initial breaking contact cluster for the three main unfolding classes “a2/b2” (orange), “a5” (purple) and “a3” (red). The breakpoints of all contact clusters are also shown as vertical lines. The unfolding curves are examples of unfolding with a short intermediate, with a longer intermediate, and an almost instantaneous unfolding without intermediate. **E** Relative unfolding curves of secondary structure elements. The percentages of broken contacts of a secondary structure element are shown as function of the overall unfolding. Each line represents a per protein average. **F** Comparison of the relative unfolding of select secondary structure elements as function of the initial unfolding class. The relative unfolding curves (E) are summarized by their area-under-thecurve (AUC); larger AUC values indicate an earlier relative unfolding and error bars the standard deviation between trajectories. **G** Distances in native protein structures of the contact cluster between a2 and b2 that initiates unfolding in “a2/b2” as function of the per protein fraction of simulations that unfold via “a2/b2”. Shown are mean and standard deviation of the different contacts within this contact cluster. Larger distances generally define less stable contacts. **H** The number of native contacts in the a5 helix and then length of the C-terminus as function of the likelihood of unfolding via the “a5”.

Small proteins can disintegrate almost instantaneously irrespective of where they were first destabilized. To test whether the different initial breakpoints actually lead to different unfolding pathways, I computed relative unfolding curves for each secondary structure that quantify how many of their contacts have been broken as function of the overall unfolding. Clear differences were readily visible for representative profiles based on per protein averages (Figure 3E). Specifically, in most proteins b2 and b3 unfold early, as visible from the fast rising curves (Figure 3E). However, in some proteins b2 and b3 unfold only at a later stage as visible from curves that initially stay flat and only rise later (Figure 3E). Remarkably, the opposite could be observed for b5 and b6 (Figure 3E). These findings suggested two major unfolding pathways, from the N- and the C-terminus, respectively.

To test whether these differences were a consequence of the initial unfolding, I next investigated the relative unfolding curves (Figure 3E) as function of the unfolding class (Figure 3C). Using the area-under-the-curve (AUC) as summary characteristic of the relative unfolding profiles, larger AUC values indicated earlier stage, while lower AUC values later stage unfolding. Importantly, averaging over trajectories by unfolding class “a1” and “a2/b2” systematically revealed an earlier unfolding of b2 and b3 compared to b5 and b6, while initial breakpoints in “a3”, “a4/b5”, and “a5” linked to the reverse order of unfolding b5 and b6 first (Figure 3F). As a further control to determine that the unfolding pathway was directly linked to the initial breakpoint, I also examined the unfolding of the *α*-helices. Trajectories of unfolding class “a1” indeed saw a1 as the first helix to unfold, as indicated by the highest AUC value (Figure 3F), and similarly for a3 in “a3”, a4 in “a4/b5” and a5 in “a5”. Thus, the initial breakpoints directly influenced the unfolding pathways that partitioned into unfolding from the N-terminal or C-terminal half. Of note, experimental evidence exists for alternative folding pathways via the N- and C-terminus in bacterial homologues [44].

Given that individual proteins had clear preferences for unfolding classes and pathways (Figures 3C,D), I sought to understand how this was encoded in the protein structures. Strikingly, the native distances of the characteristic contact cluster that breaks first in “a2/b2” increased with the propensity to unfold via “a2/b2” (Figure 3G). Longer contact distances are generally less stable, thus indicating a selective weakening. Conversely, proteins that unfolded via “a5” where characterized by a destabilizing reduction of native contacts in a5 (Figure 3H) as well as shorter C-terminal regions that may otherwise protect and anchor the C-terminus from unfolding (Figure 3H). While proteins routinely unfold stochastically from different sites, it appeared that here unfolding happened via local structural weaknesses that were by design, which in the engineering world is referred to as predetermined breaking point. Taken together, these findings suggest structure-encoded predetermined breaking points that initiate specific unfolding pathways.

### Edge cases highlight the importance of context and overlapping constraints

Why do these proteins unfold via different preprogrammed pathways? A common consequence of the unravelling of folded proteins is a heightened risk of misfolding and aggregation [45]. To estimate any change in aggregation propensity upon unfolding, I computed surface aggregation scores [46] for each frame along the unfolding simulation trajectories (Figures 4A,B). Remarkably, the thus observed increases in surface aggregation propensities directly correlated with progression of unfolding and exposure of previously buried sequences (Figure 4B). Moreover, while all proteins exhibited similar distributions of aggregation scores in their folded states (Figure 4C), the increases in aggregation propensity upon unfolding differed markedly (Figure 4C). However, there was no detectable link between an increase in aggregation propensity and how these proteins unfolded. The one exception characterised by substantially higher aggregation scores in both the folded and unfolded states was SRP102 (YKL154W). SRP102 functions differently from the other GTPases in this dataset in that it is a subunit of the signal recognition particle, a protein-RNA complex that facilitates the translocation of proteins from the cytosol to the endoplasmic reticulum [47].

**Fig. 4.**
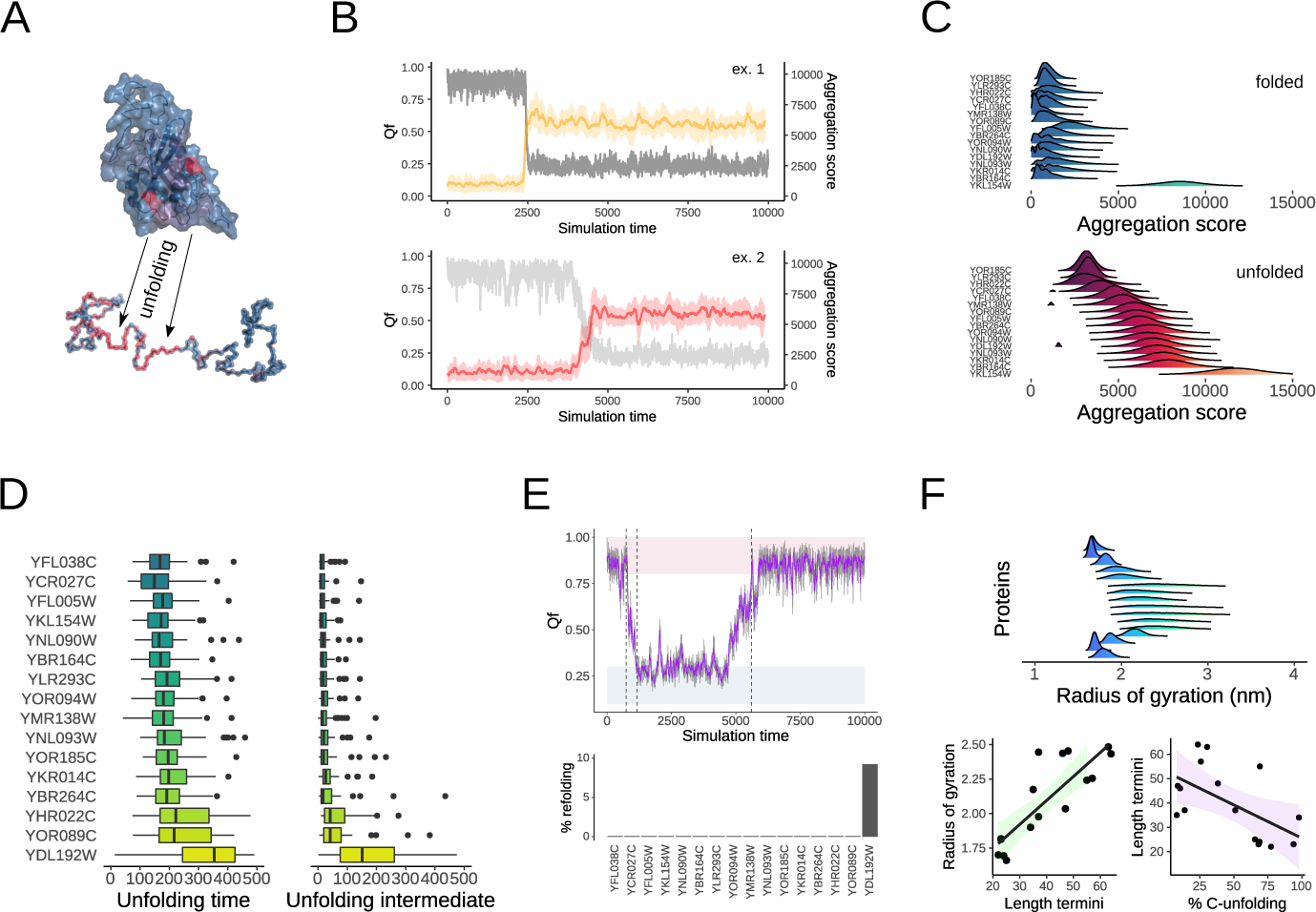
Overlapping constraints and context in complex folding landscapes. **A** Illustration of protein surface aggregation scores in folded and unfolded states. Two aggregation hotspots (red) from the folded state come in closer proximity in the unfolded state thereby much further increasing the aggregation propensity. **B** Exemplary unfolding trajectories show the time evolution of *Q_f_* and the surface aggregation score *S_agg_*. **C** Distributions of surface aggregation propensities in folded and unfolded states. **D** Distributions of unfolding times and intermediates inferred from the simulations. Time is in simulation steps. **E** An exemplary simulation trajectory of ARF1 (YDL192W) exhibits unfolding and spontaneous refolding. ARF1 is the only protein that spontaneously refolds in ca. 10% of the simulations given the current simulation length. **F** Distributions of the radius of gyration of the folded states. Differences trace to the length of the termini, especially the C-terminus. Importantly, the length of the termini correlates similarly with C-unfolding.

Similarly, some trajectories unfolded almost instantaneously while others populated intermediates (Figure 3D). The unfolding times inferred from the simulations correlated directly with the presence and length of unfolding intermediates (Figure 4D). Again, one protein, ARF1 (YDL192W) stood out from the rest by much longer unfolding times and intermediates. Remarkably, ARF1 also had by far the lowest number of native contacts and folding temperature (Figure 1C). Moreover, ARF1 was the only protein for which spontaneous refolding could be observed during the length of the simulations (Figure 4E). Beyond this, no link between unfolding pathway and unfolding rate was apparent.

Last, I observed clear differences in the compactness of the proteins’ folded states as quantified by their radius of gyration (Figure 4F), which could however almost completely be explained by the length of the termini (Figure 4F), especially the C-terminus. A higher likelihood to unfold via the C-terminus, i.e. with initial breakpoints in “a3”, “a4/b5”, and “a5”, was accompanied by shorter termini (Figure 4F), and specifically shorter C-termini (Figure 3H). Taken together, extreme cases highlight the importance of context, while overlapping constraints make protein folding landscapes rather complex.

### Predetermined breaking points are depleted in degrons

Thus far, yeast GTPases were found to differentially unfold along pathways initiated by structure-encoded local weaknesses akin to predetermined breaking points. However, the advantages of this were not clear. To test whether interaction with the protein homeostasis network influenced how the proteins unfolded, I first evaluated the presence or absence of degrons near the predetermined breaking points. Degrons are recognition sequences for ubiquitin-dependent degradation that are usually protected by the native structures [48, 49]. Strikingly, degrons were depleted in a5 in proteins that unfolded via their C-terminus (Figure 5A). Importantly, this observation was stronger when only considering highly probable degrons, but equally detectable when computing overall average degron probabilities (Figure 5A). Thus, degradation signals were systematically depleted from the vicinity of C-terminal predetermined breaking points.

**Fig. 5.**
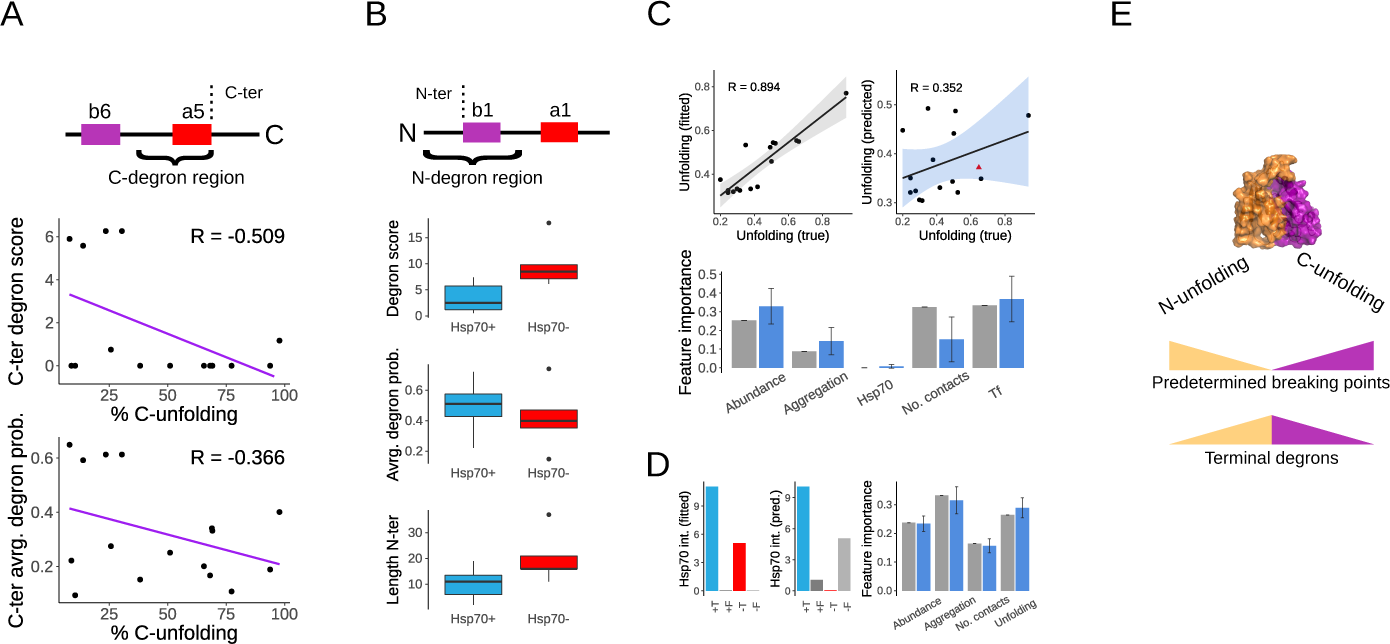
Coordination of protein (un)folding and exposure of protein homeostasis interaction signals. **A** Schematic of the C-terminal degron region. The degron score as sum over occurrences of highly probable degron sites, as well as the average degron probability are shown as function of the per protein fraction of simulation trajectories that unfold via their C-terminus. **B** Schematic of the N-terminal degron region. Distributions of the degron score, average degron probability and N-terminus length are shown for substrates (Hsp70+) and non-substrates (Hsp70-) of the cotranslationally acting Hsp70 chaperone Ssb. **C** Prediction of unfolding rates from protein and cellular features. Fitted values used the full data for training and prediction, while predicted values are based on leave-oneout cross-validated training/prediction. Feature importances from Random Forest regression models indicate the contributions of individual features. **D** Predictions of true and false (T/F) Hsp70 Ssb interactions (+/-) with Random Forst classifiers are shown as in (C) together with corresponding feature importances. **E** Summarizing schematic. *S.cerevisiae* small GTPases unfold via the N- or C-terminal in a programmed manner along predetermined breaking points that are depleted of degron degradation signals.

In contrast, a depletion of degrons from the N-terminal region of proteins that unfold via their N-terminus could be still be detected but was much weaker (R = -0.16). Because the contranslationally acting Hsp70 chaperones Ssb1,2 also bind to N-terminal hydrophobic stretches [32, 33], I sought to test whether chaperone binding and degrons were overlapping or excluding. Surprisingly, the proteins that do interact with Ssb showed a much lower degron score but a higher average degron probability (Figure 5B). This translated to heightened overall N-terminal hydrophobicity but lower degron specificity in Ssb substrates. In addition, the lower degron score in chaperone substrates was amplified by shorter N-termini (Figure 5B). These results suggested that Ssb binding and degrons ultimately rely on different specificities, while their shared hydrophobic nature possibly masked an observable depletion of N-terminal degrons in N-unfolding proteins.

Finally, I used machine learning to ask whether some the observed aspects of heterogeneous folding landscapes could be rationalized in more complex models of protein and cellular features. Herein, model fitting gave an initial indication of whether the response variable was explainable by the input features at all, while cross-validated predictions revealed the strength of more generalisable trends. The average unfolding time from the simulations (Figure 4D) could be fitted by Random Forest regression to the input features of protein abundance, aggregation score, chaperone interaction, number of native contacts and folding temperature (Figure 5C). Surprisingly, this model retained some predictive power upon cross-validation (Figure 5C) with the caveat that this was only valid when omitting the protein VPS21 (YOR089C) from the training set. The corresponding feature importances revealed that abundance, aggregation propensity and folding temperature *T_f_* increased in weight upon cross-validation, while the number of native contacts became less important and chaperone interaction played no role in trying to rationalize the unfolding times (Figure 5C). I also tried to predict interactions with the Hsp70 chaperone Ssb from some of the newly derived unfolding features. Importantly, while Hsp70 interactions could be fitted with Random Forst classifiers to abundance, unfolded surface aggregation score, native contacts and unfolding time, there was no discriminative power, and an uninformative mix of feature importances after cross-validation in this small dataset (Figure 5D). In fact, the only signal that correlated with Hsp70 Ssb interaction was a higher average degron probability, i.e. higher sequenced hydrophobicity in the N-terminal region (Figure 5B). These results are consistent with the previous finding in a larger dataset that the by far strongest determinator of Hsp70 Ssb interactions in small single-domain proteins seems to be the presence of recognition sites as opposed to characteristics of the folding landscape [32]. While the present dataset of 16 proteins is too small to learn all the complex relationships encoded within proteins, it is noteworthy that several other observations such as the propensity of N- or C-unfolding could not even be fitted to global protein or cellular characteristics. Instead, they appeared to exclusively depend on the local encoding of the initial breaking points.

## Discussion

This work revealed how proteins from the same organisms and with near identical structures can display systematically heterogeneous folding landscapes. Yeast GTPases (un)folded either via the N- or C-terminus according to an initial loss of stability in structure-encoded predetermined breaking points. In turn, degrons as recognition signals for ubiquitin-dependent degradation were depleted from the initial unfolding sites, as if to protect the protein from too rapid degradation upon spontaneous unfolding (Figure 5E).

Proteins are highly dynamic, and frequently unfold and refold locally. Degrons are generally effective, as shown for example by destabilizing mutations that increase degron exposure and thus degradation [50]. It is possible to speculate that strong degrons that are usually buried should only become accessible upon complete unfolding or misfolding, not immediately after local unfolding or already before the folding is completed, especially in proteins that spontaneously unfold and refold during their lifecycle. Similarly, the finding that degrons and Hsp70 Ssb binding encode different specificities should make sense in this context. There is no a priori competition at the ribosome between chaperones and ubiquitin ligases for nascent chains [51]. Rather, a large number of proteins in yeast interacts cotranslationally with Ssb [33] while cotranslational degradation remains the rare exception for a fraction of the particularly folding challenged proteins [52].

The main caveat with the presented results lies in their level of resolution. The use of coarse-grained structure-based models that only consider native contacts made this work computationally feasible, but ignored the role of non-native contacts in protein misfolding and folding intermediates. Similarly, unfolding simulations are computationally efficient but only serve as a proxy for the folding pathways. Fortunately, in both cases the underlying assumptions and simplifications are generally assumed valid for small single-domain proteins. Moreover, the analysis of protein unfolding is of central interest in its own right, as presented here, because it is the precursor of regulated protein turn-over, condensate formation, or, alternatively, misfolding and aggregation. To this end, individual small GTPases have been studied in more detail at the atomistic level, and further work will be required to fully map out their folding landscapes. However, any comparative analyses depends on a level of resolution that serves as a common denominator. The hope is that the presented results will motivate future studies to characterize *in vivo* protein folding landscapes include the role of the protein termini [53], and the interplay between translation, folding, and protein export pathways [54].

Finally, accumulating evidence suggests that the protein homeostasis network acts as much as a central regulator of cell functionality through controlling levels of functional proteins than as a garbage collector for damage proteins. For instance, significant subsets of both chaperones and ubiquitin ligases were found to systematically coexpress with genes involved in synapse formation and maintenance in the human brain [55]. Similarly, the temperature sensitivity of embryonic development was found to depend on protein homeostasis [56]. Protein sensors based on temperature-sensitive local unfolding [57] are one example of protein homeostasis feedback cascades that may act far beyond the well characterized heat-shock response [58]. These ideas also readily apply to yeast small GTPases whose immediate functions lie protein transport, which however if disrupted, for instance by not being folded properly, can bring about broad consequences for the regulation of cell signalling [59]. Thus, there are many reasons for continuing to seek to understand *how* proteins fold. Especially, understanding how proteins dynamically fold and unfold in tight interaction with their cellular environment will likely hold the keys to unravelling much of their currently not understood complexity.

## Methods

### Data and computer code availability

All project data and computer code to reproduce the presented results is available at https:/www.github.com/pechmannlab/UNfold.

### Data sources

The set of carefully equilibrated protein structures and structural models of 16 homologous *S.cerevisiae* small GTPases, as well as their structure-based multiple sequence alignment were obtained from previous work [32].

### Molecular modelling

Equilibrium protein dynamics were probed with all-atom explicit solvent molecular dynamics (MD) simulations performed with GROMACS [60]. Following a standard protocol of energy minimization as well as 1*ns* NVT and 1*ns* NPT equilibration, three independent 10*ns* MD production runs were generated for each protein.

Unfolding simulations were performed as *C_α_*-coarse-grained structure-based models [42, 61] at the protein’s folding temperature *T_f_*. The folding temperature *T_f_* was determined through a raster search of short simulations at different temperatures and identification of the temperature corresponding to the system’s maximum heat capacity by the weighted histogram method [62]. For each protein, 70 independent unfolding simulations of 10*ns* (2 *·* 10^7^ simulation steps, 10^4^ reported time frames) were run. Only trajectories that included an unfolding event were included in further analyses; the median number of trajectories per protein included in further analyses was 53, the first quartile 46, and the minimum 29. All simulations were prepared with the sbmOpenMM Python library [63] and run in openMM [64]. The trajectories were subsequently analysed with MDTraj [65].

### Data analysis

Secondary structure was assigned with DSSP [66], and consensus secondary structure elements were defined through regions of agreeing secondary structure assignments across the alignment. The numbers of native contacts were obtained from the *C_α_* structure based models. The all encompassing set of native contacts from all proteins was mapped onto the alignment and hierarchically clustered by amino acids (AA) distance, i.e. proximity along the backbone. For example, two contacts were recorded at the alignment positions 45-76, and 46-77. The AA distances between these two contacts was now computed as the sum of the distances between the residues at positions 45-46 and 76-77, in each case upon identifying the corresponding amino acids in the individual proteins, measuring their distances on the protein structures, and finally averaging over all proteins. Clusters of contacts with a minimum separation of 1*nm* were defined as contact clusters that contained on average 5 individual contacts.

Frustration in protein structures was computed with FrustratometeR [67]. The map of correlated motions was derived from the distances profiles of the contact clusters. Specifically, the profiles of contact distances along the MD trajectories were averaged to obtain average distance profiles for each contact cluster. Moreover, the average profiles of three independent MD runs were concatenated. Correlated motions were then detected by Pearson’s correlation coefficients *R >* 0.5 between these profiles.

To describe protein unfolding, two related metrics were used. The order parameter *Q_f_*as the fraction of formed contacts considered all contacts within 120% of their initial distances as ‘formed’, if exceeding as ‘broken’. *Q_f_*is particularly helpful to describe the overall unfolding along a trajectory. In contrast, the breakpoints of the contact clusters were defined based on distance profiles smoothed by averages over a sliding window of size *w* = 20 and a hard threshold of 2.5*nm*. Specifically, many contacts initially break and reform frequently, and a smoothed profile denoises this a little. Moreover, the threshold-based breakpoint, i.e. the simulation time frame when the distance between a pair of residues exceeded 2.5*nm* during the unfolding window, was detected for each individual contact within a contact cluster. Next, the largest cluster of most similar breakpoints was identified by hierarchical clustering of the different breakpoint within a contact cluster, and its median breakpoint recorded as the contact cluster breakpoint. Of note, in almost all cases the breakpoints within a contact cluster were practically identical. The additional steps of clustering and consensus building only served to filter out rare outliers that originated from using hard thresholds on noisy data. The unfolding class was assigned based on a majority vote of the most represented location among the first five breaking contact clusters in a simulation trajectory.

Sequence protein aggregation propensities were predicted with Tango [40], and surface aggregation scores [46] computed as 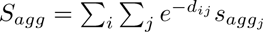. Herein, the sum extends over all solvent exposed amino acids *i* for which a surface patch aggregation score is computed as the sum over the sequence aggregation scores *s_agg_j* of itself (*i* = *j*) and all neighbouring solvent exposed residues *j* weighted by their distances *d_ij_* in an exponential decay term. I.e., close aggregation prone neighbours contribute more, and distant residues do not contribute noticeably. Solvent exposure was determined with the function “shrake rupley” of the Python library MDTraj and a threshold of 0.7.

Random Forest regression and classification models were fitted with the Python library sklearn. Yeast protein abundances were obtained from the PaxDB database [68]. Degrons were predicted with the software provided by [49], and consensus Hsp70 Ssb interactions obtained from [32]. Protein structures were visualized with PyMOL (Schrödinger, LLC).

## Declarations

The author declares no conflicts of interest.

